# Complexities in the role of acetylation dynamics in modifying inducible gene activation parameters

**DOI:** 10.1101/2021.05.17.444476

**Authors:** Samantha Carrera, Amanda O’Donnell, Yaoyong Li, Karol Nowicki-Osuch, Syed Murtuza Baker, David Spiller, Andrew D. Sharrocks

**Affiliations:** Faculty of Biology, Medicine and Health, University of Manchester, Michael Smith Building, Oxford Road, Manchester, M13 9PT, UK

**Keywords:** Acetylation, deacetylation, EGF signaling, transcription

## Abstract

High levels of histone acetylation are associated with the regulatory elements of active genes, suggesting a link between acetylation and gene activation. However, several studies have shown that histone acetylation dynamics rather than hyperacetylation per se are important determinants in gene activation, particularly at inducible genes. We revisited this model, in the context of EGF-inducible gene expression and found that rather than a simple unifying model, there are two broad classes of genes; one in which high lysine acetylation activity is required for efficient gene activation, and a second group where the opposite occurs and high acetylation activity is inhibitory. We examined the latter class in more detail using *EGR2* as a model gene and found that lysine acetylation levels are critical for several activation parameters, including the timing of expression onset, and overall amplitudes of the transcriptional response. In contrast, *DUSP1* responds in the canonical manner and its transcriptional activity is promoted by acetylation. Single cell approaches demonstrate heterogenous *DUSP1* activation kinetics and that acetylation levels influence allele activation frequencies. Our data therefore point to a complex interplay between acetylation dynamics and target gene induction, which cannot simply be explained by a unified response to acetylation activity. Instead, acetylation level thresholds are an important determinant of transcriptional induction dynamics that are sensed in a gene-specific manner.

## Introduction

Histone modifications play an important role in controlling the levels of gene transcription. High levels of histone acetylation are associated with transcriptionally active genes where the modifications decorate the nucleosomes surrounding both proximal and distal regulatory elements (Schubeler et al., 2004; Roh et al., 2004; reviewed in Anderson and Sandelin, 2020). It is generally assumed that high levels of histone acetylation are associated with active transcription and reciprocally, low levels or hypoacetylation are associated with repressed or inactive genes. However, evidence for a more dynamic model has been gathered, chiefly through the use of histone acetylation/deacetylation inhibitors, whereby histone acetylation dynamics rather than overall levels are critical in gene activation (Hazzalin and Mahadevan, 2005; Crump et al., 2011; reviewed in Clayton et al., 2006). It is important to note that in addition to histones, other transcriptional regulatory proteins can be targeted by acetylation to influence their activity as exemplified by studies on MMTV promoter activation (Mulholland et al., 2003). Thus, acetylation dynamics may have an impact beyond simply through modifying histones associated with regulatory elements.

Lysine acetylation levels are governed by the combined actions of lysine acetyl transferases (KATs) and lysine deacetylases (KDACs) found at gene regulatory elements. By using acute administration of KAT (Crump et al., 2011) and KDAC (Hazzalin and Mahadevan, 2005) inhibitors, both types of enzymatic activity have been implicated as important for the inducible activation of genes such as *FOS* and *JUN*, providing weight to the model that acetylation dynamics rather than overall levels are critical for inducible transcription to take place. Furthermore, genome-wide ChIP-seq studies have also shown a role for both KDACs and KATs at active genes providing further support for a role for dynamic histone acetylation in gene activation (Wang et al., 2009). This has led to the broader recognition that KDACs and their deacetylation activity can have activating roles in addition to their previously established role in transcriptional repression (reviewed in Smith, 2007). For example, class I KDACs are required for the inducible activation of glucocorticoid receptor target genes (Kadiyala et al., 2013). As both KATs and KDACs are considered as therapeutically targets in a range of human diseases (reviewed in Brown et al., 2016; Falkenberg and Johnstone, 2014), it is therefore important to further understand their mechanisms of action.

Many of the insights provided to date have focused on a relatively small number of genes and, in particular, the growth factor regulated *FOS* and *JUN*. We therefore wished to more broadly assess the role of acetylation dynamics in modifying the activation of growth factor inducible genes. Rather than a simple unifying model, we identified two broad classes of genes; one in which high acetylation activity is required for efficient gene activation, and a second group where the opposite occurs and high acetylation is inhibitory. This points to a more complex model where acetylation levels thresholds are an important determinant of transcriptional induction dynamics that are sensed in a gene-specific manner.

## Results

### EGF-inducible gene expression profiles in MCF10A cells

To establish the repertoire of EGF-inducible genes in MCF10A cells, we treated cells with EGF and used RNA-seq to profile mRNA expression. We first generated RNA-seq data at two time points, namely 0 and 30 minutes to identify genes showing rapid induction. A total of 212 genes exhibited enhanced expression at 30 minutes (1.5 fold; adjusted p-value <0.05 Supplementary Table 1) (Nowicki-Osuch, et al., 2017). To further interrogate the expression profiles of these genes, we generated another RNA-seq dataset at 4 time points over a 3 hour time period (Fig. 1A). There are several clearly distinct expression profiles, with a group of genes exhibiting rapid and transient responses while others show more delayed and/or extended responses to EGF stimulation (Fig. 1A). To study the mechanisms of induction further we decided to focus on two rapidly induced genes, *EGR2* and *DUSP1. EGR2*, shows a robust transient response to EGF stimulation and *DUSP1* shows lower amplitude response (see Fig. 2B). To allow real-time continuous monitoring of gene induction kinetics, we used CRISPR-Cas9 editing to create MCF10A reporter lines which contain a fusion of the nanoluciferase gene (Hall et al., 2012) to one allele of either *EGR2* or *DUSP1* (Fig. 1B; Supplementary Fig. 1A and B). The nanoluciferase has been engineered by incorporating PEST sequences to create a short half life of 10-30 mins thus enabling its use as a readout for monitoring transient induction kinetics. Importantly, the induction kinetics of the *EGR2-luciferase* fused allele closely mimicked that of the endogenous transcript (Supplementary Fig. 1C). Indeed, in keeping with the RNA-seq analysis, the *EGR2* reporter exhibited the expected high amplitude and transient luciferase activity kinetics whereas the *DUSP1* reporter showed more extended activity (Fig. 1C). Importantly, both reporter genes showed reduced induction following treatment with the MEK inhibitor trametinib, demonstrating the expected ERK-dependent signalling response (Fig. 1C).

**Fig. 1.**
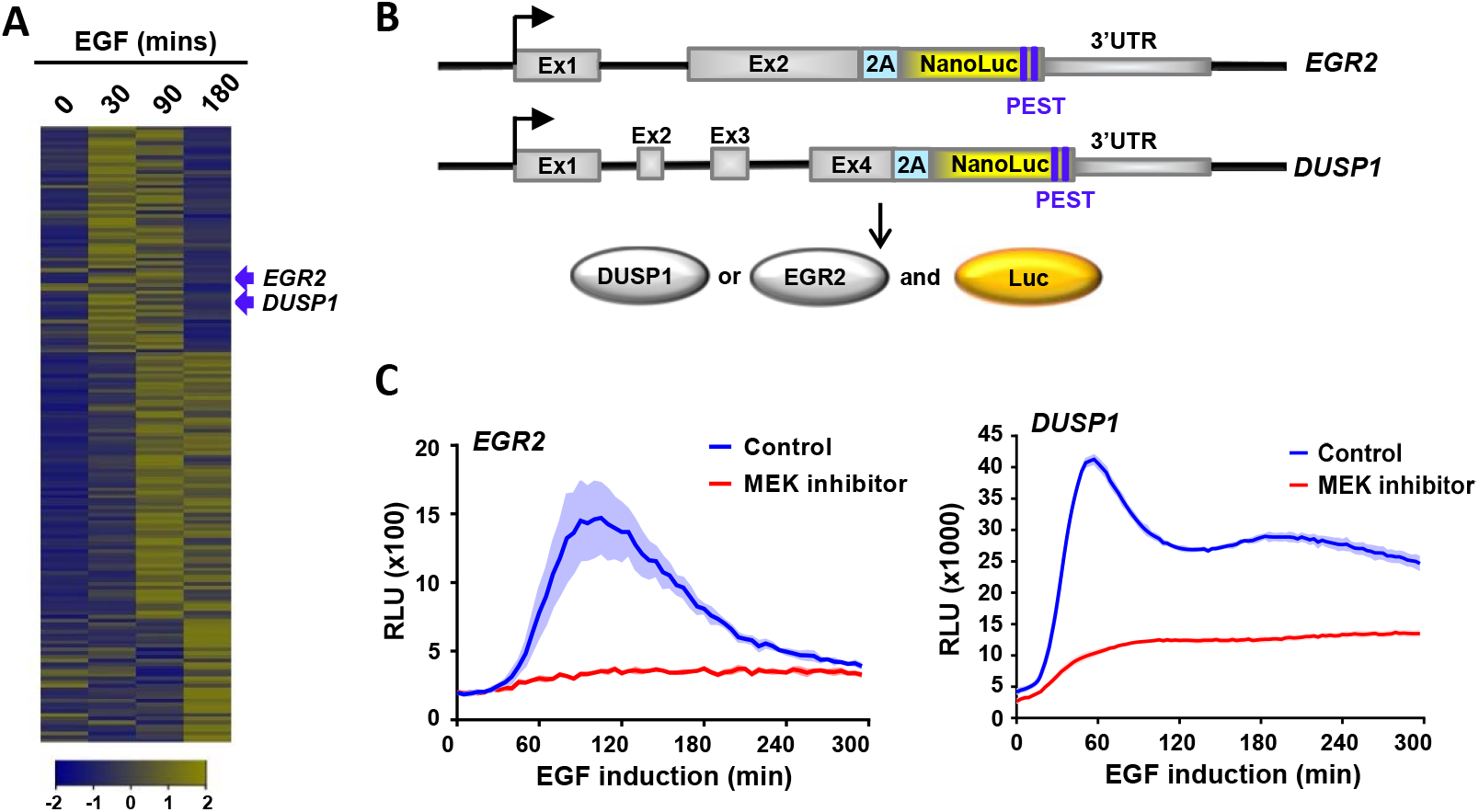
Establishment of a reporter gene assay to study EGF-mediated gene activation kinetics. (A) RNA-seq analysis of EGF inducible genes in MCF10A cells at the indicated time points following EGF addition. Data are shown for genes with FPKM >0.1 as row z-normalised values. (B) Diagrammatic illustration of the luciferase reporter constructs where nanoluciferase was introduced into the 3’ end of genomic *EGR2* or *DUSP1* loci in MCF10A cells. There is a 2A sequence inserted after their coding regions to allow expression of non-fused nanoluciferase. A PEST sequence is incorporated to allow for rapid degradation. (C) Activation kinetics of *EGR2*- and *DUSP1*-luciferase reporters after induction with EGF. Cells were treated with vehicle or the MEK inhibitor trametinib 1 hr prior to EGF addition. Data are the average of three biological replicates (n=3), shaded area represents ±SEM.

**Fig. 2.**
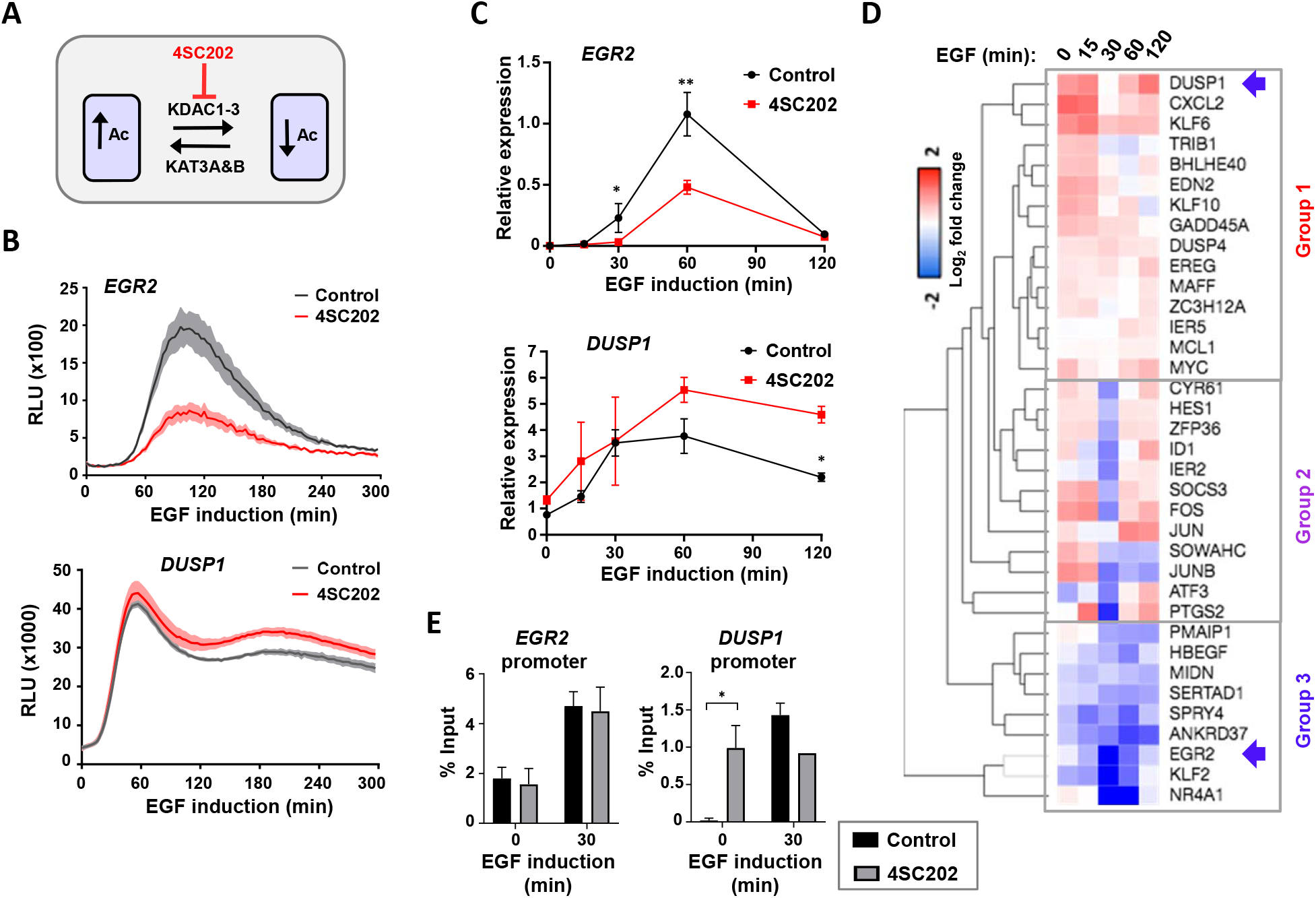
Differential effects of KDAC inhibitors on EGF-mediated gene activation. (A) Model showing the point of action of the class I HDAC inhibitor GNE-781 in the acetylation-deacetylation equilibrium. (B) Activation kinetics of *EGR2*- and *DUSP1*-luciferase reporters after induction with EGF. Cells were treated with vehicle or 4SC202 1 hr prior to EGF addition. Data are the average of three biological replicates (n=3), shaded area represents ±SEM. (C and D) RT-qPCR analysis of relative mRNA expression profiles of the indicated genes after induction with EGF for the indicated times. Cells were treated with vehicle (Control) or 4SC202 for 1h before induction. Detailed profiles for *EGR2* and *DUSP1* are shown in (C). Data represents mean of three biological replicates ± SD; *= P-value < 0.05, **= P-value <0.001. The heatmap in (D) is from RT-qPCR analysis using the Fluidigm Biomark system and shows log_2_ fold changes of gene expression caused by 4SC202 administration after EGF induction at the indicated times. Data are the average of three biological replicates (n=3). Samples are grouped according to similar response profiles to KDAC inhibition. (E) H3K27ac levels at the *EGR2* or *DUSP1* promoters after treatment with 4SC202 for 1h prior to adding EGF for 30 min. Data represents the mean of three biological replicates ± SD; *= P-value < 0.05.

### Complex changes to the EGF induction profiles following deacetylase inhibition

Having established two reporter systems, we next examined their response to changes in acetylation activity. First, we used the specific class I lysine deacetylase inhibitor 4SC202 (Pinkerneil, et al. 2016) to block deacetylation activity (Fig. 2A), and administered this shortly before (60 mins) EGF induction to monitor acute responses. The *DUSP1* reporter showed increased activity following treatment with 4SC202 as would be expected by inhibiting deacetylases, which are thought to be repressive in nature (Fig. 2B, bottom). However, in contrast, *EGR2* reporter activity was severely curtailed by the deacetylase inhibitor (Fig. 2B, top). This differential response to deacetylase inhibition was unexpected but this was verified at the mRNA level by RT-qPCR (Fig. 2C; Supplementary Fig. S2A). This effect appears specific to EGF inducible genes as acute administration of deacetylase inhibitors has little effect on the expression of non-inducible *HADHB* control gene (Supplementary Fig. S2A). Furthermore, an alternative deacetylase inhibitor, sodium butyrate, caused similar effects on EGF-inducible gene expression, extinguishing induction of the *EGR2* reporter, but enhancing *DUSP1* reporter activity (Supplementary Fig. S2B). Given these divergent responses, we widened our analysis to a panel of 36 EGF inducible genes and performed RT-qPCR analysis over an induction time course. Three different responses to deacetylase inhibition were observed; group 1 (including *DUSP1*) generally had increased expression across the time course consistent with transcriptional derepression; group 2 showed reduced peak expression, whereas group 3 (including *EGR2*) showed broadly reduced expression across the time course (Fig. 2D; Supplementary Fig. S3). Similar effects were seen at both the mature RNA and pre-mRNA levels as exemplified by *SOCS3, CXCL2* and *EREG*, demonstrating an effect on nascent transcription levels (Supplementary Fig. S3).

To explore the potential effects of deacetylase inhibition on the histone acetylation levels found at gene regulatory elements, we monitored H3K27 acetylation levels at the *EGR2* and *DUSP1* promoters. As expected, significant increases were observed to basal acetylation levels at the *DUSP1* promoter following HDAC inhibition (Fig. 2E). EGF stimulation enhanced acetylation at this promoter but no further increase was elicited after HDAC inhibition (Fig. 2E). However, deacetylase inhibition has no effect on the steady state acetylation levels at the *EGR2* promoter (Fig. 2E). Only the molecular changes at *DUSP1* promoter are therefore consistent with the effects of HDAC inhibition on its transcriptional output. However, for *EGR2*, a simple direct cause and effect relationship on histone acetylation levels at its promoter is hard to establish.

Collectively these data demonstrate that there is a heterogenous effect of inhibition of class I deacetylases on EGF-mediated gene expression. Individual genes show either enhanced or reduced expression following deacetylase inhibition, rather than just the general increases expected from inhibiting deacetylase-mediated transcriptional repression.

### Acetylation inhibitors cause reciprocal responses to EGF-mediated gene induction profiles

Given the unexpected complexities in response to deacetylase inhibition, we asked whether acute acetylase inhibition causes reciprocal effects on the EGF target gene induction profiles. We focussed on two inhibitors of P300/CBP (KAT3B/KAT3A), A485 which is thought act broadly on histone acetylation levels at promoter and enhancer elements (Lasko, et al., 2017), and GNE-781 which is thought to predominantly act at enhancer elements (Fig. 3A; Romero et al., 2017; Raisner et al., 2017). The *DUSP1* reporter activity profile was dampened down after treatment with the A485 inhibitor and also by administration of the GNE-781 inhibitor at the early stages of induction (Fig. 3B). These effects on *DUSP1* transcription are consistent with the known association between high levels of histone acetylation and gene activation. In contrast however, the *EGR2* reporter activity is elevated following treatment with either acetylation inhibitor (Fig. 3B). These effects on reporter activity were broadly recapitulated by directly measuring *EGR2* and *DUSP1* mRNA levels (Fig. 3C). Thus, reciprocal effects are seen on both *EGR2* and *DUSP1* expression kinetics following inhibition of acetylation and deacetylation, with high acetylation activity being associated with increased activation of *DUSP1* but decreased activation of *EGR2*. Importantly, acute administration of acetylase inhibitors has little effect on the expression of non-inducible control genes such as *GLUD1* and *HADHB* (Supplementary Fig. S5A) which is consistent with what we observed with deactetylase inhibitors.

**Fig. 3.**
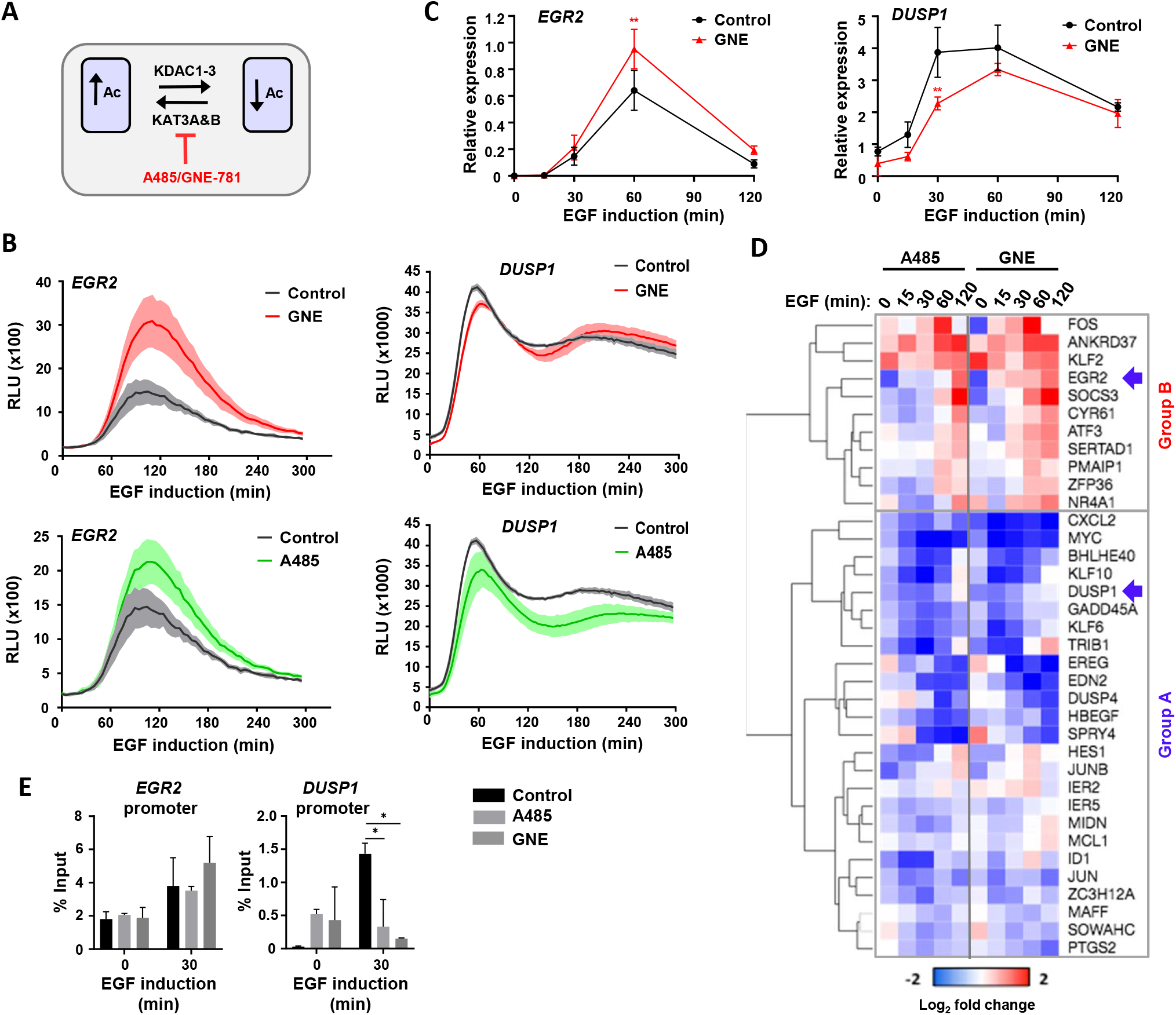
Differential effects of KAT inhibitors on EGF-mediated gene activation. (A) Model showing the point of action of the KAT3A/B inhibitors A485 and GNE-781 in the acetylation-deacetylation equilibrium. (B) Activation kinetics of *EGR2*- and *DUSP1*-luciferase reporters after induction with EGF. Cells were pre-treated with vehicle (Control), A485 or GNE-781 prior to EGF addition. Data are the average of three biological replicates (n=3), shaded area represents ±SEM. (C and D). RT-qPCR analysis of relative mRNA expression profiles of the indicated genes after induction with EGF for the indicated times. Cells were treated with vehicle (Control) or GNE-781 1 hr prior to EGF addition. Detailed profiles for *EGR2* and *DUSP1* are shown in (C). Data represents mean of three biological replicates ± SD; *= P-value < 0.05, **= P-value <0.001. The heatmap in (D) is from RT-qPCR analysis using the Fluidigm Biomark system and shows log_2_ fold changes of gene expression caused by KAT inhibitor administration after EGF induction at the indicated times. Data are the average of three biological replicates (n=3). Samples are grouped according to similar response profiles to KAT inhibition. (E) ChIP-qPCR analysis of H3K27ac levels at the *EGR2* or *DUSP1* promoters after induction with EGF for 30 min after prior treatment with vehicle (Control), A485 or GNE-781. Data represents mean of three biological replicates ± SD; *= P-value < 0.05, **= P-value <0.001.

We next examined whether this reciprocality is observed more generally across the EGF-activated programme, and found two broad categories of genes; group B which is generally upregulated by inhibiting acetylation (including *EGR2*) and a larger group (A) whose expression is reduced (including *DUSP1*) (Fig. 3D). There is generally a high level of consistency in the response to the two inhibitors. Similar effects were seen at both the mature RNA and pre-mRNA levels as exemplified by *SOCS3, CXCL2* and *EREG*, demonstrating an effect on nascent transcription levels (Supplementary Fig. S5B and C). Reduced acetylation levels would not be expected to result in enhanced transcriptional output, and we therefore asked whether these are the same genes identified as decreasing in activity following deacetylase inhibition. All of the genes showing enhanced expression following acetylation inhibition (group B) are in the two groups (2 and 3) which show either decreased expression at all, or a subset, of time points following deacetylase inhibition (compare Fig. 2D and Fig. 3D; Supplementary Fig. S4A). This reinforces the view that decreased acetylation levels lead to the enhanced expression of a cohort of genes (group B) in response to EGF induction. This differential response to disturbing the acetylation equilibrium, suggests that group A genes may share different properties to group B genes, and examination of their expression profiles demonstrates that group B genes generally show earlier peak induction and a faster decrease back towards basal activity (Supplementary Fig. S4B).

Next, we asked whether changes in H3K27 acetylation could be found at the *EGR2* and *DUSP1* regulatory elements which could explain the transcriptional regulatory effects we observed. No changes could be observed in H3K27 acetylation at the *EGR2* promoter following acetylase inhibition (Fig. 3E). In contrast, we observe attenuation of the EGF-mediated increases in *DUSP1* promoter acetylation following treatment with either acetylation inhibitor (Fig. 3E), which is fully consistent with the reduced transcription levels caused by the same treatment. Taken together with the deacetylase inhibitor results, changes to *DUSP1* promoter acetylation occur as predicted from disturbing the acetylation equilibrium and are sufficient to explain the changes in *DUSP1* expression according to established paradigms linking promoter acetylation to gene activation. However, there is no change to histone acetylation levels at the *EGR2* promoter after disrupting the acetylation-deacetylation equilibrium and therefore no obvious link to the changes in its expression.

In summary, we observe a broad degree of reciprocality in effects of manipulating the acetylation equilibrium on gene expression levels, although two different categories of inducible genes can be identified; those that show the expected decrease in expression when acetylation activity is increased and those that show the opposite behaviour to manipulating the acetylation-deacetylation equilibrium.

### The impact of acetylation activity levels on transcriptional induction parameters

Having established the impact of acetylation levels on bulk cell populations we next turned to single cell analysis using our *EGR2*-luc and *DUSP1*-luc reporter lines to further elucidate the mechanism of action. At the single cell level, the response to EGF stimulation is highly heterogeneous (Fig. 4A; Supplementary Fig. S6A; Supplementary movies 1 and 2). In the case of *EGR2*, there is negligible baseline reporter expression and very few cells exhibit high amplitude induction (>10 fold) and the majority fall below this threshold, with some registering barely detectable luciferase activity. In contrast, baseline expression of *DUSP1* is detectable in most cells, and their magnitude of induction shows a narrower range (ranging between 2.2 and 10 fold) (Supplementary Fig. 6A). To further investigate the gene induction profiles we normalised the data to remove fluctuations in gene expression amplitude and used principal component analysis (PCA) to group the different expression profiles (Supplementary Fig. S6B). This identified two characteristic profiles for *EGR2* and three different profiles for *DUSP1* (Fig. 4B; Supplementary Fig. S6C). For both genes, the timing of their peak expression is a key defining parameter, indicating a non-uniform response of cells to EGF induction.

**Fig. 4.**
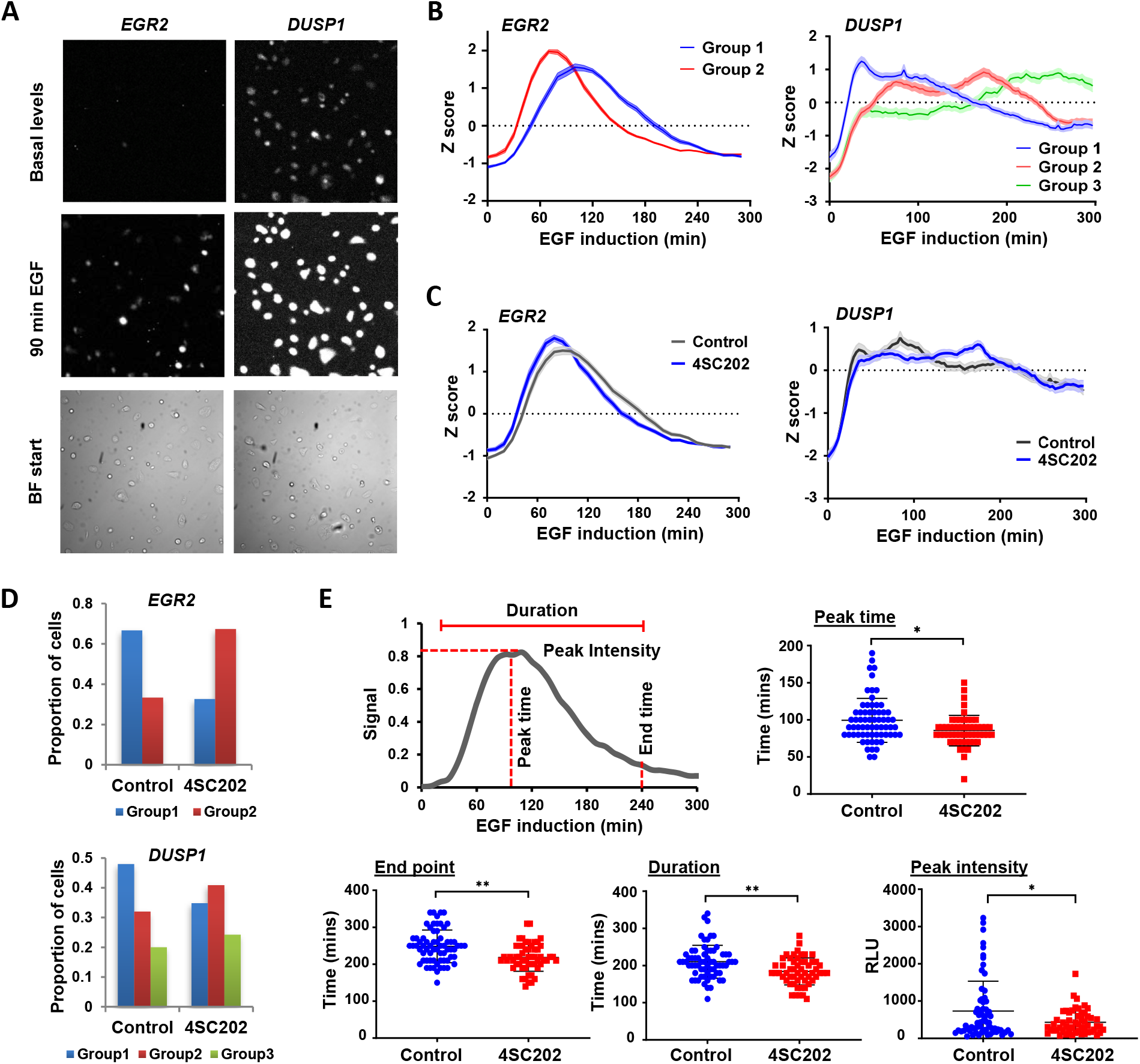
Single cell analysis of KDAC inhibition effects reveals complex changes in gene induction parameters. (A) Examples of bioluminescence microscope images of *EGR2*-(left) and *DUSP1*-(right) reporters in unstimulated cells (basal level) or in cells stimulated with EGF for 90 min. Bright field (BF) images show cells prior to induction. (B) Expression kinetics of *EGR2*-(left) and *DUSP1*-(right) reporters for the different profiles defined by PCA analysis (groups 1-3; see Supplementary Fig. S6B and C). The data includes a combination of all single cells measured (*EGR2* n=109; *DUSP1* n=100). Lines represent average Z scores of profiles from single cells ±SEM (shaded area). (C) Kinetics of expression of *EGR2*- and *DUSP1*-reporters after either treatment with vehicle or 4SC202 for 1h prior to EGF addition. The bioluminescence of single cells was measured and the lines represent the average Z score of profiles from single cells. The shaded area represents ±SEM. (D) Ratio of cells exhibiting EGF-mediated *EGR2*- or *DUSP1*-reporter expression profiles belonging to the groups identified by PCA analysis (groups 1-3; see Supplementary Fig. S6B and C) after treatment with vehicle (Control) or 4SC202. (E) The schematic shows different parameters of *EGR2*-reporter activity measured in single cells following EGF addition. These parameters are plotted with (n=60) and without (n=49) 4SC202 addition. In the graphs, each dot represents one cell. *= P-value < 0.05, **= P-value <0.001.

Next, we inhibited deacetylation and asked whether this impacted on various activation parameters. To study activation kinetics more carefully, we normalised for differences in the amplitude of activation, and found that the expression profile of *EGR2* is shifted, indicating earlier initiation (Fig. 4C). For *DUSP1*, the changes appear more complex but the initial phase of activation follows the same trajectory in the presence or absence of inhibitor (Fig. 4C). To investigate these changes more closely we returned to the different single cell profiles identified by PCA analysis. For *EGR2*, we still see two distinct profiles but there is a shift in the numbers of cells exhibiting the group 1 pattern to group 2, thereby favouring an earlier response to EGF (Fig. 4D). For *DUSP1*, we see more complex changes to the expression profiles with a loss of group 1, compensated for by a gain in group 2 and 3 profiles. We further examined various parameters associated with the *EGR2* expression profile, including peak time, end point, duration and peak intensity and found statistically significant changes in all parameters at the single cell level when treated with deacetylase inhibitor (Fig.4E). The peak time and the end point are shifted earlier, consistent with the normalised data (Fig. 4C). There is also a decrease in the duration of response and the overall peak intensity following deacetylase inhibition, emphasising a multifaceted response to manipulating acetylation activity.

To provide further insights into EGF-inducible gene activation kinetics, we turned to smFISH to examine the effects on allelic activation frequencies following deacetylase inhibition. First we analysed the distribution of total number of mRNA molecules across different cells. The results were broadly in agreement with the luciferase reporter assays with *EGR2* exhibiting a broader distribution of mRNA molecules per cell upon induction in comparison to the tighter distribution associated with *DUSP1* (Supplementary Fig. S7). Treatment with the deacetylase inhibitor 4SC202 caused a general reduction in the numbers of *EGR2* transcripts per cell whereas the opposite was observed for *DUSP1* (Supplementary Fig. S7), consistent with what we observed using reporter alleles or bulk mRNA from cell populations. To study allelic activation frequencies, we focussed on *DUSP1*. In the absence of EGF stimulation very few cells contained active transcriptional loci but after 15 mins stimulation, over 60% of cells showed activation of 2 or 3 alleles (MCF10A cells contain an extra copy of the chromosomal segment harbouring the *DUSP1* locus) (Fig. 5A and B). Following that, there is a general decline to one or no active alleles by 60 mins. However, in the presence of the deacetylase inhibitor 4SC202, a large number of cells already show activation of one or more alleles under basal conditions and there is a clear increase in the proportion of cells containing 2 or 3 active alleles after 15 mins EGF treatment which is sustained at 30 mins. In contrast when cells were treated with an acetylation inhibitor, only a small proportion of cells show activation of 2 or 3 alleles at any timepoint after EGF stimulation. Thus, acetylation-mediated control over the number of active alleles likely plays a major role in determining the outcome of transcription at the *DUSP1* locus.

**Fig. 5.**
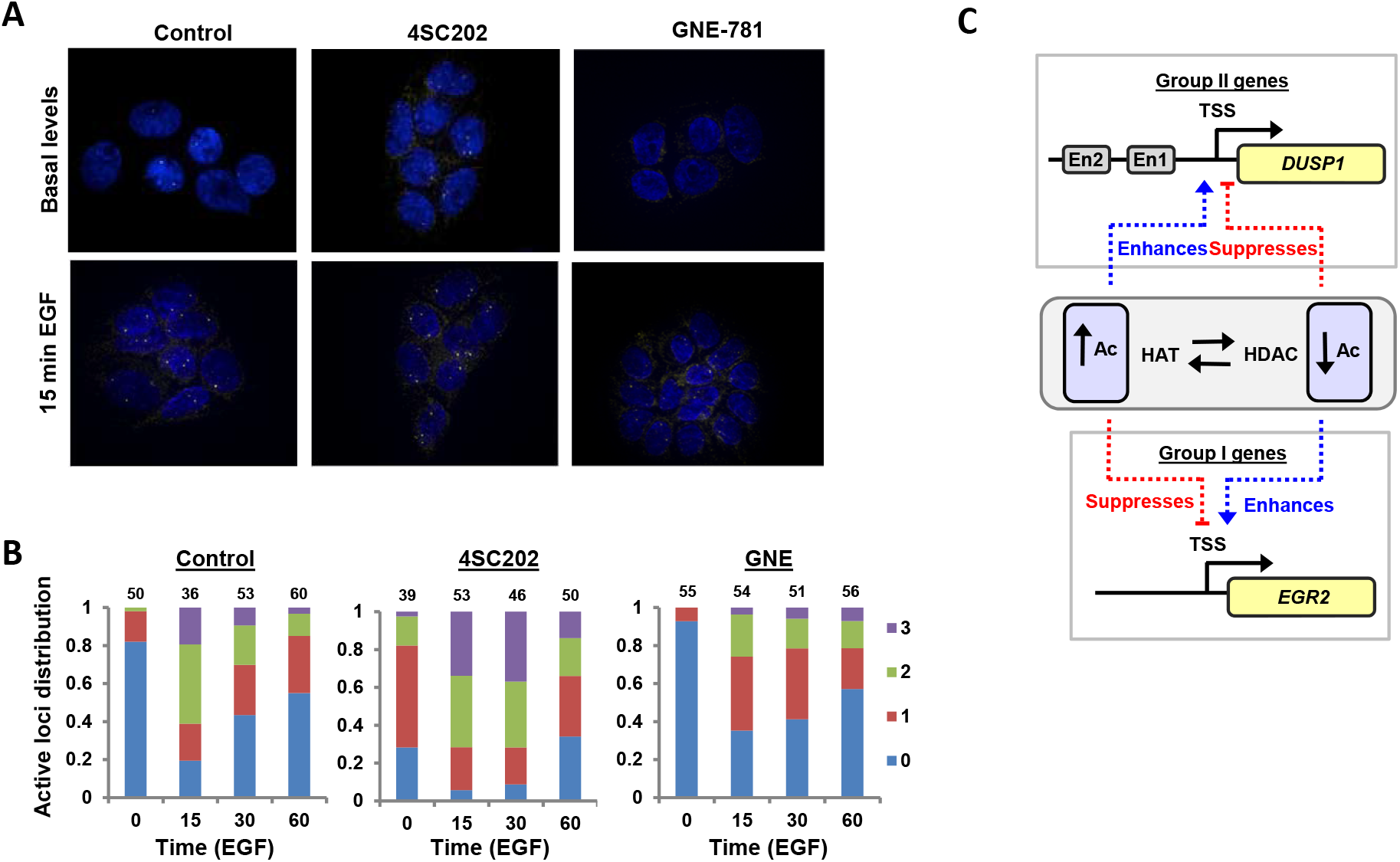
*DUSP1* allele activation frequency in response to disturbing the acetylation-deacetylation equilibrium. (A and B) smFISH of *DUSP1* mRNA production in serum starved (basal) and EGF stimulated MCF10A cells in the presence of prior addition of vehicle, 4SC202 or GNE-781. Representative images from cells following 15 mins induction are shown in (A). (B) Quantification of the distribution of cells with 0, 1, 2 or 3 active transcription sites at each time point and condition. Number at the top of each column indicates number of cells (n). (C) Model showing the effects of altering the acetylation (Ac) equilibrium on the activation of the EGF inducible genes *DUSP1* and *EGR2*, examples of larger groups of genes that respond in a similar manner. The transcriptional start site (TSS) and putative enhancer regions (En) are indicated.

Collectively, these single cell approaches reveal a complex series of gene activation parameters that are altered by changing acetylation dynamics that underly the expression profiles derived from cell populations.

## Discussion

High levels of histone acetylation are typically associated with transcriptionally active genes both at promoter and enhancer elements and this has led to the assumption that increased acetylation leads to high level gene transcription as exemplified by the use of KAT inhibitors on a genome-wide scale (Raisner et al., 2018; Weinert et al., 2018). Nevertheless, reciprocal effects are observed at some loci such as genes encoding core regulatory transcription factors where KDAC inhibitors suppress transcription (Gryder et al., 2019). This indicates that high level acetylation does not always lead to enhanced transcriptional activity. Here we examined the effect of disturbing the acetylation equilibrium catalyzed by class I KDACs and the KATs P300/CBP on growth factor inducible gene activation. We found that EGF activated genes fall into two broad categories (Fig. 5C); those which show the canonical activation response to increased acetylation levels (group A eg *DUSP1*) and those which show the opposite effect and whose expression is dampened when acetylation levels are increased (group B e.g. *EGR2*). Thus, rather than a unified response, different genes are tuned to respond differently to changes in acetylation levels, even when responding to the same signaling events.

Using single cell approaches, we gained further insights into the gene activation dynamics at *DUSP1* and *EGR2* and the mechanistic changes induced by disturbing the acetylation equilibrium. Rather than a uniform response to EGF addition, there is a large variation in the magnitude of peak *EGR2* transcription, but there are two basic expression profiles which differ according to their timing onset. In contrast, the *DUSP1* expression profile is much more complex, with a more uniform induction level across single cells but three different underlying kinetic profiles. KDAC inhibition led to a switch in the timing of *EGR2* expression to the earlier of the two kinetic profiles and a concomitant drop in overall peak expression levels and the duration of activation. The *DUSP1* activation profiles were also altered by KDAC inhibition leading to a shift towards later expression onset and peak expression. However, due to the more complex expression profiles for *DUSP1*, we were unable to further disentangle the changes elicited by KDAC inhibition. Instead we examined allelic activation frequencies and found that KDAC inhibition increased the numbers of active alleles per cell whereas KAT inhibition had the opposite effect. Perturbing the acetylation dynamics therefore affects the allele activation frequency. This is reminiscent of recent studies where KAT inhibition reduced *Fos* transcriptional burst frequencies in neurons (Chen et al., 2019) and, reciprocally, high KAT activity and high histone acetylation levels augmented burst frequencies of *Bmal1* and hence transcriptional output (Nicolas et al., 2018).

The effects of disturbing the acetylation equilibrium on transcriptional levels can be explained largely by the changes to H3K27ac at the *DUSP1* promoter, and a recent study demonstrated that a subset of genes respond to acetylation changes at their promoters (Hsu et al., 2021). However, the latter study made the surprising finding that H3K27ac acetylation levels at most promoters are refractory to P300/CBP inhibition and here we show that *EGR2* falls into this category. The acetylation event(s) that are altered at the *EGR2* locus therefore remain enigmatic. In many ways, our findings that the effects of KDAC inhibitors on *EGR2* activation are independent of obvious changes to promoter histone H3K27 acetylation are reminiscent of findings on MMTV transcription where the effects of KDAC inhibitors are attributed to changes to the basal transcription machinery rather than to histones (Mulholland et al., 2003). This suggests a broader effect of KDACs in gene activation beyond their role in histone modifications. However, it should be noted that there are numerous different histone modifications in addition to H3K27ac that we tested here, and it is possible that KDACs function on one or more of these in the context of enhancing EGF-mediated genes activation. Others have questioned the relevance of H3K27ac in activation of regulatory elements and their associated genes and found that this mark is generally not needed for enhancer activity in mouse ESCs (Zhang et al., 2020). It is possible that other acetylation marks are more relevant, including others such as H3K18ac H3K56ac, H3K64ac, and H3K122ac that are deposited by P300/CBP (Das et al., 2009; Di Cerbo et al., 2014; Tropberger et al., 2013). Moreover, H4K5/8/12ac have been shown to be important at promoters for signal-dependent transcriptional activation (Hargreaves et al., 2009) and their levels may be influenced by KDAC inhibition. Alternatively, there may be other non-histone proteins involved whose activity is influenced by acetylation levels (Weinert et al., 2018).

Overall, we identified a group of genes (group A: exemplified by *DUSP1*) whose activity is affected as expected by disturbing acetylation levels, with high acetylation causing increased levels of gene transcription but another set of genes (group B: exemplified by *EGR2*) behaves in the opposite manner. Interestingly, at this second group of genes, acetylation inhibitors show an effect earlier in the induction profile than deacetylation inhibitors (see Supplementary Fig. S4). This suggests a model whereby initial high level basal acetylation caused by KDAC inhibition is refractory for the onset of gene activation. Conversely, the inducible acetylation changes that occur following growth factor stimulation are more affected by the acetylation inhibitors where the dampening down of acetylation levels leads to a transcriptional overshoot. This implies that high acetylation levels normally limit the overall activation levels of expression at this set of genes.

While the growth factor inducible genes fall into these two categories based on KAT inhibition profiles, they respond subtly differently to KDAC inhibition, with group B genes (activated by KAT inhibition) being split between two subgroups, one of which shows general dampening across the profiles and another where only the peak expression is reduced. Thus, although there are commonalities, there are further subtle differences in how individual genes respond, likely caused by their unique regulatory setups and different induction profiles. Indeed, previous work on *FOS* induction, demonstrated the importance of the acetylation-deacetylation equilibrium for gene activation, and in that case inhibition of both KDACs and KATs reduced transcriptional levels (Hazzalin and Mahedevan, 2005; Crump et al., 2011). In combination with these earlier studies, our data therefore indicate that it is important to re-evaluate the assumptions that steady state acetylation levels mean that acetylation promotes transcriptional activity. Instead, this should be evaluated on a case by case basis, where each gene is tuned to respond differently to dynamically changing acetylation levels catalyzed by the opposing functions of KDACs and KATs.

## Supporting information

Supplementary Table S2- primers

Supplementary Table S1- EGF regulated gene list

Movie showing single cell EGR2-luc activation

Movie showing single cell DUSP1-luc activation

## Acknowledgements

We thank Guanhua Yan and Mairi Challinor for excellent technical assistance; Peter March and Steve Marsden in the Bioimaging facility and Mike White for bioimaging advice; Claire Morrisroe in the Genomic Technologies facility. We also thank Nicoletta Bobola, Shen-His Yang and members of our laboratory for comments on the manuscript and stimulating discussions. This work was funded by the Wellcome Trust (103857/Z/14/Z). The Bioimaging Facility SMC microscopes used in this study were purchased with grants from BBSRC, Wellcome Trust and the University of Manchester Strategic Fund.

## Author contributions

The conception or design of the work (SC, ADS), the acquisition, analysis, or interpretation of data (SC, DS, SMB, KNO), drafting the work or revising it critically (SC, KNO, ADS). All authors approved the manuscript and are accountable for all aspects of the work.

## Materials and methods

### Cell line creation and culture

Parental MCF10A cells and cell lines derived from these were grown in DMEM/F12 (Gibco, 11320–033) containing 5% horse serum (Biosera, DH291), 20 ng/ml EGF (Sigma, E1257), 10 μg/ml insulin (Sigma, I0516), 100 ng/ml cholera toxin (Sigma, C9903) and 0.5 μg/ml hydrocortisone (Sigma, H0396) (complete medium). When required cells were seeded or changed into starvation media: DMEM/F12 containing 0.5% horse serum, 10 μg/ml insulin, 100 ng/ml cholera toxin and 0.5 μg/ml hydrocortisone. Cells were left for 48h in starving media and then treated with EGF where required.

To create MCF10A-EGR2-Luc cells, we used the pX335 CRISPR/Cas9 plasmid (Cong et al 2013; obtained via Addgene) and the *EGR2* 3’-end-targetting guide RNA sequences; target 1 (ADS5212, ADS5213) and target 2 (ADS5214, ADS5215). These guides were inserted into plasmid pX335 BbsI cloning site as described previously (Ran et al. 2013) to create plasmids pX335-EGR2-target1 (pAS4883) and pX335-EGR2-target2 (pAS4884). For donor plasmid EGR2-2A-LucP (pAS4885), the full-length nanoLuciferasePEST gene (pNL1.2 Promega) was PCR-amplified along with a 63 bp 5’-end viral 2A sequence using primers ADS5225/ADS6756 then prepared for Gibson assembly using primers ADS5227/ADS5228. The luciferase gene was then cloned into the BamHI site in the pUC19 backbone vector using the Gibson assembly technique, along with the PCR-amplified 782 bp left homology arm (primers ADS5220/ADS5226 incorporating a single base mutation to remove PAM) and 876 bp right homology arm (primers ADS5229/ADS5230), from genomic DNA for specific integration by homologous recombination. Low passage MCF10A cells were transfected at 80% confluency in a 25 cm^2^ flask using 25 μl FugeneHD (Promega) with 6 μg donor plasmid (pAS4885), 1 μg each CAS9 guide RNA plasmid (pAS4883 and 4884) and 200 ng linear hygromycin resistance gene (Clontech) were used. Transfected cells were selected for by adding hygromycin (50 μg/ml) for 72 hours before plating surviving cells onto 10 cm dishes without antibiotic selection. Single cells were seeded in 96-well plates using serial dilutions, inserts were checked using primers ADS5337 and ADS5338.

To create MCF10A-DUSP1-Luc cells, we used AIO-GFP as a source of Cas9-D10A (Chiang, et al. 2016; obtained via Addgene) and the *DUSP1* targeting guide RNA sequences (ADS6055, ADS6056, ADS6057, ADS6058) were inserted into the AIO-GFP plasmid to create pAS4863. A HDR template containing the full-length nanoLuciferasePEST gene (pNL1.2 Promega) and *DUSP1* homology arms was synthesised as a double-stranded DNA GBlock (Integrated DNA Technologies). Low passage MCF10A cells were transfected at 80% confluency in a 25cm^2^ flask using 25 μl FugeneHD (Promega) with 6 μg of pAS4863 and 200 ng of GBlock donor template. Transfected cells were grown for 72 hours and then cell sorted into a GFP positive population. GFP positive cells were grown for further 5 days before seeding single cells in 96-well plates using serial dilutions, inserts were checked using primers ADS6061 and ADS6141. All of the PCR primers used are detailed in Supplementary Fig. S2.

### RNA extraction and qRT-PCR

RNA was extracted using the RNeasy plus kit (Qiagen) following the manufacturer’s protocol. Subsequently, RNA samples were quantified using a Nanodrop 2000 (Thermo Scientific) and concentrations were normalized to 20 ng/μl. 40 ng of each sample was used per reverse transcription-quantitative polymerase chain reaction (RT-qPCR) reaction using the QuantiTect SYBR^®^ Green RT-PCR Kit (Qiagen, 204243) on Rotor-Gene Q (Qiagen) real-time PCR machine. When indicated, the nanolitre volume RT-qPCR was performed using the Fluidigm Biomark HD system using EvaGreen chemistry following the manufacturer’s protocol. Six housekeeping genes were included in the analysis (*GLUD1, C3, HADHB, SRSF6, GAPDH* and *HPRT1*). The geometric mean of the housekeeping genes was calculated for each sample. Samples were normalized to the mean of the housekeeping genes before calculating DeltaCT. The PCR primers are detailed in Supplementary Fig. S2. The output data were processed following the default quality protocol. Data points with more than one peak in the melt analysis were discarded.

### RNA-seq library preparation

The nuclear RNA-seq data were generated from the EGF-induced MCF10A cells at two time points, 0 minute and 30 minutes as previously described (E-MTAB-5370; Nowicki-Osuch et al., 2017). Additionally, RNA-seq was performed on whole cell RNA extracts using longer EGF time course (0, 30, 90, 180 min) as previously described (Nowicki-Osuch et al., 2017). Briefly, MCF10A cells were seeded in medium without EGF and with 0.5% Horse Serum 48 hours prior to EGF stimulation. EGF (final concentration 20 ng/ml) was added for indicated times and RNA was extracted using an RNeasy plus kit (Qiagen, 74134) with DNase treatment according to the manufacturer’s protocol. Samples with RNA integrity number (RIN) > 9 were used for sequencing library construction with the TruSeq Stranded mRNA sample preparation protocol (Illumina).

### RNA-seq analysis

Cufflinks was used to identify differential expression (DE) genes between the two time points 0 min and 30 min (Trapnell, et al., 2012). The R package edgeR was also used to identify DE genes between 0 min and 30 min (Robinson et al., 2010). In order to obtain the EGF inducible genes (i.e. those genes with significantly higher transcription level at 30 minutes in comparison with that at 0 minute), the same criteria, namely fold change >1.5 and adjusted p-value <0.05, was applied separately on the DE genes obtained from Cufflinks and edgeR, resulting in 168 genes from Cufflinks and 160 genes from edgeR. We merged together the two sets of genes and obtained 212 unique EGF inducible genes (Supplementary table 1). Among the 212 genes in the list, 169 genes were selected whose mean FPKM is >0.1 in a new data set of nuclear RNA-seq at four time points of EGF induction (E-MTAB-5370) and were used to draw the heatmap plot in Fig. 1A.

### Chromatin Immunoprecipitation

MCF10A cells (4×10^6^) were seeded in starvation media 48 hours before induction with EGF. Cells were induced with 20 ng/ul EGF for the desired time. Cells were cross linked for 10 minutes in 1% Formaldehyde before quenching with 0.125 M Glycine for 5 min. Cells were harvested in ice-cold PBS with complete protease inhibitor (Roche) and washed sequentially with ChIP Lysis Buffer I (50 mM Hepes-KOH pH 7.5, 140 mM NaCl, 1 mM EDTA, 10% Glycerol, 0.5% Igepal, 0.25% Triton X-100) and ChIP Lysis Buffer II (10 mM Tris-HCl pH 8, 200 mM NaCl, 1 mM EDTA pH 8, 0.5 mM EGTA pH 8.0). The resulting nuclei were resuspended in ChIP Lysis Buffer III (10 mM Tris-HCl pH 8.0, 100 mM NaCl, 1 mM EDTA pH 8.0, 0.5 mM EGTA pH 8.0, 0.1% Na-Deoxycholate, 0.5% N-lauroylsarcosine). Lysates were sonicated on ice to yield 200-600 bp DNA fragments using a Bioruptor (Diagenode).

Magnetic protein A Dynabeads (Invitrogen) were incubated with 0.5 μg of anti-H3K27ac antibody (Abcam - ab4729) or non-specific IgG overnight at 4°C. Antibody/beads were washed and incubated with nuclear extracts overnight at 4°C. Immunoprecipitates were washed 5 times with RIPA buffer (50 mM HEPES-KOH pH 7.6, 500 mM LiCl, 1 mM EDTA pH 8.0, 1% Igepal, 0.7% Na-Deoxycholate) and once with TE-NaCl (10mM Tris-HCl pH 8.0, 1mM EDTA, 50 mM NaCl). ChIP DNA was eluted from Protein A Dynabeads by adding 150 μl elution buffer (50 mM Tris-HCl pH 8.0, 10 mM EDTA pH 8.0, 1% SDS) and incubating at 65°C. Cross-links were reversed by heating to 65°C overnight, then treating with proteinase K for 1 h at 45°C. Chromatin was cleaned using QiaQuick PCR cleanup columns (Qiagen). Nanolitre volume qPCR was performed using the Fluidigm Biomark HD system using EvaGreen chemistry following manufacturer’s protocol and the PCR primers detailed in Supplementary Fig. S2. The output data were processed following the default quality protocol. Data points with more than one peak in the melt analysis were discarded.

### Bioluminescence detection in live cells

For analyzing cell population timecourses, MCF10A-EGR2-Luc or MCF10A-DUSP1-Luc cells were seeded into 96-well plates in starvation media 48 hours before the experiment. Where required, cells were treated with 1.2 nM 4SC202 (Selleckchem), 100 nM A485 (Selleckchem), 100 nM GNE-781 (Romero et al., 2017; kindly supplied by Karen Gascoigne, Genentech) or vehicle (DMSO) for one hour prior to induction with EGF. NanoGlo Endurazine substrate (Promega) was added to each well 30 min before EGF induction. Cells were induced with 20 ng/ml EGF (Sigma) and luminescence was measured for 5 seconds every 5 min in FLUOstar Omega microplate reader.

For single cell analysis, MCF10A-EGR2-Luc or MCF10A-DUSP1-Luc cells were seeded into a 35/10 MM glass bottom dish (Greiner Bio-One) in starvation media 48 hours before the experiment. NanoGlo Endurazine substrate (Promega) was added to each well 30 min before EGF induction. Where required, cells were treated with 1.2 nM 4SC202 or vehicle (DMSO) for one hour prior to induction with EGF. Images were collected on a Zeiss Axio Observer A.1 microscope using a *10x / 0*.*5 Fluar* objective and an iXon Ultra EMCCD Camera (Andor) camera through Micromanager software v1.4.15. Images were then processed and analysed using Fiji ImageJ (http://imagej.net/Fiji/Downloads).

To identify similar patterns in single cell profiles, principal component analysis (PCA) was conducted on normalised counts of luminescent signals from *DUSP1*- and *EGR2*-*luciferase* transgenes across the seven time-points. UMAP visualization was then generated by taking the first 10 principal components for *DUSP1* and 15 principal components for *EGR2*.

### smFISH

smFISH probes for *EGR2* and *DUSP1* were designed and ordered using the Stellaris Probe Designer (Biosearch Technologies). MCF10A cells were seeded into glass coverslips in starving media and left to grow for 48 hours before inducing them with EGF and treated with inhibitors where required. Media was removed and cells were fixed with formaldehyde solution for 15 min at room temperature, washed and incubated in 70% ethanol for 2 hours. Samples were hybridized using 100 nM Stellaris probes for *EGR2*, or *DUSP1* overnight. Coverslips were mounted using ProLong Gold antifade Mountant with DAPI (ThermoFisher). Images were acquired on an Olympus IX83 inverted microscope using Blue, Red and Green-Yellow Lumencor LED excitation, a *60x/ 1*.*42 Plan Apo* objective and the *Sedat* filter set (Chroma *89000*). The images were collected using a *R6 (Qimaging)* CCD camera with a Z optical spacing of [0.2μm]. Raw images were then deconvolved using the Huygens Pro software (SVI) and *maximum intensity* projections of these deconvolved images are shown in the results. Mature mRNA transcripts and transcription site quantification was performed using FISHquant (Mueller, et al., 2013).

### Statistical analysis

Data for qRT-PCR and ChIP-qPCR are presented as means of a minimum of three biological replicates. Two-way ANOVA was performed using GraphPad PRISM v8, statistical significance was determined using the Sidak multiple comparisons test. Luminometer data presented is the average ±SEM of three biological replicates with at least three technical repeats each.

Single cell luciferase data and smFISH data are the combination of at least two biological replicates, statistical significance was calculated using unpaired t test GraphPad PRISM v8.

## References

Andersson R, Sandelin A. Determinants of enhancer and promoter activities of regulatory elements. Nat Rev Genet. 2020 21(2):71–87.

Brown JA, Bourke E, Eriksson LA, Kerin MJ. Targeting cancer using KAT inhibitors to mimic lethal knockouts. Biochem Soc Trans. 2016 44(4):979–86.

Chen LF, Lin YT, Gallegos DA, Hazlett MF, Gómez-Schiavon M, Yang MG, Kalmeta B, Zhou AS, Holtzman L, Gersbach CA, Grandl J, Buchler NE, West AE. Enhancer Histone Acetylation Modulates Transcriptional Bursting Dynamics of Neuronal Activity-Inducible Genes. Cell Rep. 2019 26(5):1174-1188.e5.

Chiang TW, le Sage C, Larrieu D, Demir M, Jackson SP. CRISPR-Cas9(D10A) nickase-based genotypic and phenotypic screening to enhance genome editing. Sci Rep. 2016 6:24356.

Clayton AL, Hazzalin CA, Mahadevan LC. Enhanced histone acetylation and transcription: a dynamic perspective. Mol Cell. 2006 23(3):289–96.

Cong L, Ran FA, Cox D, Lin S, Barretto R, Habib N, Hsu PD, Wu X, Jiang W, Marraffini LA, Zhang F. Multiplex genome engineering using CRISPR/Cas systems. Science. 2013;339(6121):819–823.

Crump NT, Hazzalin CA, Bowers EM, Alani RM, Cole PA, Mahadevan LC. Dynamic acetylation of all lysine-4 trimethylated histone H3 is evolutionarily conserved and mediated by p300/CBP. Proc Natl Acad Sci U S A. 2011 108(19):7814–9

Das C, Lucia MS, Hansen KC, Tyler JK. CBP/p300-mediated acetylation of histone H3 on lysine 56. Nature. 2009 459(7243):113–7.

Di Cerbo V, Mohn F, Ryan DP, Montellier E, Kacem S, Tropberger P, Kallis E, Holzner M, Hoerner L, Feldmann A, Richter FM, Bannister AJ, Mittler G, Michaelis J, Khochbin S, Feil R, Schuebeler D, Owen-Hughes T, Daujat S, Schneider R. Acetylation of histone H3 at lysine 64 regulates nucleosome dynamics and facilitates transcription. Elife. 2014 3:e01632.

Falkenberg, K. J., Johnstone, R. W. Histone deacetylases and their inhibitors in cancer, neurological diseases and immune disorders. Nat. Rev. Drug Discov. 2014. 13:673–69.1

Gryder BE, Wu L, Woldemichael GM, Pomella S, Quinn TR, Park PMC, Cleveland A, Stanton BZ, Song Y, Rota R, Wiest O, Yohe ME, Shern JF, Qi J, Khan J. Chemical genomics reveals histone deacetylases are required for core regulatory transcription. Nat Commun. 2019 10(1):3004.

Hargreaves DC, Horng T, Medzhitov R. Control of inducible gene expression by signal-dependent transcriptional elongation. Cell. 2009 10138(1):129–45.

Hall MP, Unch J, Binkowski BF, Valley MP, Butler BL, Wood MG, Otto P, Zimmerman K, Vidugiris G, Machleidt T, Robers MB, Benink HA, Eggers CT, Slater MR, Meisenheimer PL, Klaubert DH, Fan F, Encell LP, Wood KV. Engineered luciferase reporter from a deep sea shrimp utilizing a novel imidazopyrazinone substrate. ACS Chem Biol. 2012 7(11):1848–57.

Hazzalin CA, Mahadevan LC. Dynamic acetylation of all lysine 4-methylated histone H3 in the mouse nucleus: analysis at c-fos and c-jun. PLoS Biol. 2005 3(12):e393.

Hsu E, Zemke NR, Berk AJ. Promoter-specific changes in initiation, elongation, and homeostasis of histone H3 acetylation during CBP/p300 inhibition. Elife. 2021 10:e63512.

Kadiyala V, Patrick NM, Mathieu W, Jaime-Frias R, Pookhao N, An L, Smith CL. Class I lysine deacetylases facilitate glucocorticoid-induced transcription. J Biol Chem. 2013 288(40):28900–12.

Lasko LM, Jakob CG, Edalji RP, Qiu W, Montgomery D, Digiammarino EL, Hansen TM, Risi RM, Frey R, Manaves V, Shaw B, Algire M, Hessler P, Lam LT, Uziel T, Faivre E, Ferguson D, Buchanan FG, Martin RL, Torrent M, Chiang GG, Karukurichi K, Langston JW, Weinert BT, Choudhary C, de Vries P, Kluge AF, Patane MA, Van Drie JH, Wang C, McElligott D, Kesicki EA, Marmorstein R, Sun C, Cole PA, Rosenberg SH, Michaelides MR, Lai A, Bromberg KD. Discovery of a selective catalytic p300/CBP inhibitor that targets lineage-specific tumours. Nature. 2017 550(7674):128–132.

Mulholland NM, Soeth E, Smith CL. Inhibition of MMTV transcription by HDAC inhibitors occurs independent of changes in chromatin remodeling and increased histone acetylation. Oncogene. 2003 22(31):4807–18.

Mueller F, Senecal A, Tantale K, Marie-Nelly H, Ly N, Collin O, Basyuk E, Bertrand E, Darzacq X, Zimmer C. FISH-quant: automatic counting of transcripts in 3D FISH images. Nature Methods 2013 10(4):277–278

Nicolas D, Zoller B, Suter DM, Naef F. Modulation of transcriptional burst frequency by histone acetylation. Proc Natl Acad Sci U S A. 2018 115(27):7153–7158

Nowicki-Osuch K, Li Y, Challinor M, Gerrard DT, Hanley NA, Sharrocks AD. EINCR1 is an EGF inducible lincRNA overexpressed in lung adenocarcinomas. PLoS One, 2017 12(7):e0181902.

Pinkerneil M, Hoffmann MJ, Kohlhof H, Schulz WA, Niegisch G. Evaluation of the Therapeutic Potential of the Novel Isotype Specific HDAC Inhibitor 4SC-202 in Urothelial Carcinoma Cell Lines. Target Oncol. 2016 11(6):783–798.

Ran FA, Hsu PD, Wright J, Agarwala V, Scott DA, Zhang F. Genome engineering using the CRISPR-Cas9 system. Nat Protoc. 2013 8(11):2281–2308

Robinson MD, McCarthy DJ, Smyth GK. edgeR: a Bioconductor package for differential expression analysis of digital gene expression data. Bioinformatics 2001 26(1):139–140.

Roh TY, Ngau WC, Cui K, Landsman D, Zhao K. High-resolution genome-wide mapping of histone modifications. Nat Biotechnol. 2004 22(8):1013–6.

Romero FA, Murray J, Lai KW, Tsui V, Albrecht BK, An L, Beresini MH, de Leon Boenig G, Bronner SM, Chan EW, Chen KX, Chen Z, Choo EF, Clagg K, Clark K, Crawford TD, Cyr P, de Almeida Nagata D, Gascoigne KE, Grogan JL, Hatzivassiliou G, Huang W, Hunsaker TL, Kaufman S, Koenig SG, Li R, Li Y, Liang X, Liao J, Liu W, Ly J, Maher J, Masui C, Merchant M, Ran Y, Taylor AM, Wai J, Wang F, Wei X, Yu D, Zhu BY, Zhu X, Magnuson S. GNE-781, A Highly Advanced Potent and Selective Bromodomain Inhibitor of Cyclic Adenosine Monophosphate Response Element Binding Protein, Binding Protein (CBP). J Med Chem. 2017 60(22):9162–9183.

Raisner R, Kharbanda S, Jin L, Jeng E, Chan E, Merchant M, Haverty PM, Bainer R, Cheung T, Arnott D, Flynn EM, Romero FA, Magnuson S, Gascoigne KE.. Enhancer Activity Requires CBP/P300 Bromodomain-Dependent Histone H3K27 Acetylation. Cell Rep. 2018 24(7):1722–1729.

Schübeler D, MacAlpine DM, Scalzo D, Wirbelauer C, Kooperberg C, van Leeuwen F, Gottschling DE, O’Neill LP, Turner BM, Delrow J, Bell SP, Groudine M. The histone modification pattern of active genes revealed through genome-wide chromatin analysis of a higher eukaryote. Genes Dev. 2004 18(11):1263–71.

Smith CL. A shifting paradigm: histone deacetylases and transcriptional activation. Bioessays. 2007 30(1):15–24

Trapnell C, Roberts A, Goff L, Pertea G, Kim D, Kelley DR, Pimentel H, Salzberg SL, Rinn JL, Pachter L. Differential gene and transcript expression analysis of RNA-seq experiments with TopHat and Cufflinks. 2012. Nat Protoc, 7:562–578.

Tropberger P, Pott S, Keller C, Kamieniarz-Gdula K, Caron M, Richter F, Li G, Mittler G, Liu ET, Bühler M, Margueron R, Schneider R. Regulation of transcription through acetylation of H3K122 on the lateral surface of the histone octamer. Cell. 2013 152(4):859–72.

Wang Z, Zang C, Cui K, Schones DE, Barski A, Peng W, Zhao K. Genome-wide mapping of HATs and HDACs reveals distinct functions in active and inactive genes. Cell. 2009 138(5):1019–31.

Weinert BT, Narita T, Satpathy S, Srinivasan B, Hansen BK, Schölz C, Hamilton WB, Zucconi BE, Wang WW, Liu WR, Brickman JM, Kesicki EA, Lai A, Bromberg KD, Cole PA, Choudhary C. Time-Resolved Analysis Reveals Rapid Dynamics and Broad Scope of the CBP/p300 Acetylome. Cell. 2018 174(1):231-244.e12.

Zhang T, Zhang Z, Dong Q, Xiong J, Zhu B. Histone H3K27 acetylation is dispensable for enhancer activity in mouse embryonic stem cells. Genome Biol. 2020 21(1):45.

