## Supplementary Figures 1-7 for "Complexities in the role of acetylation dynamics in modifying inducible gene activation parameters"

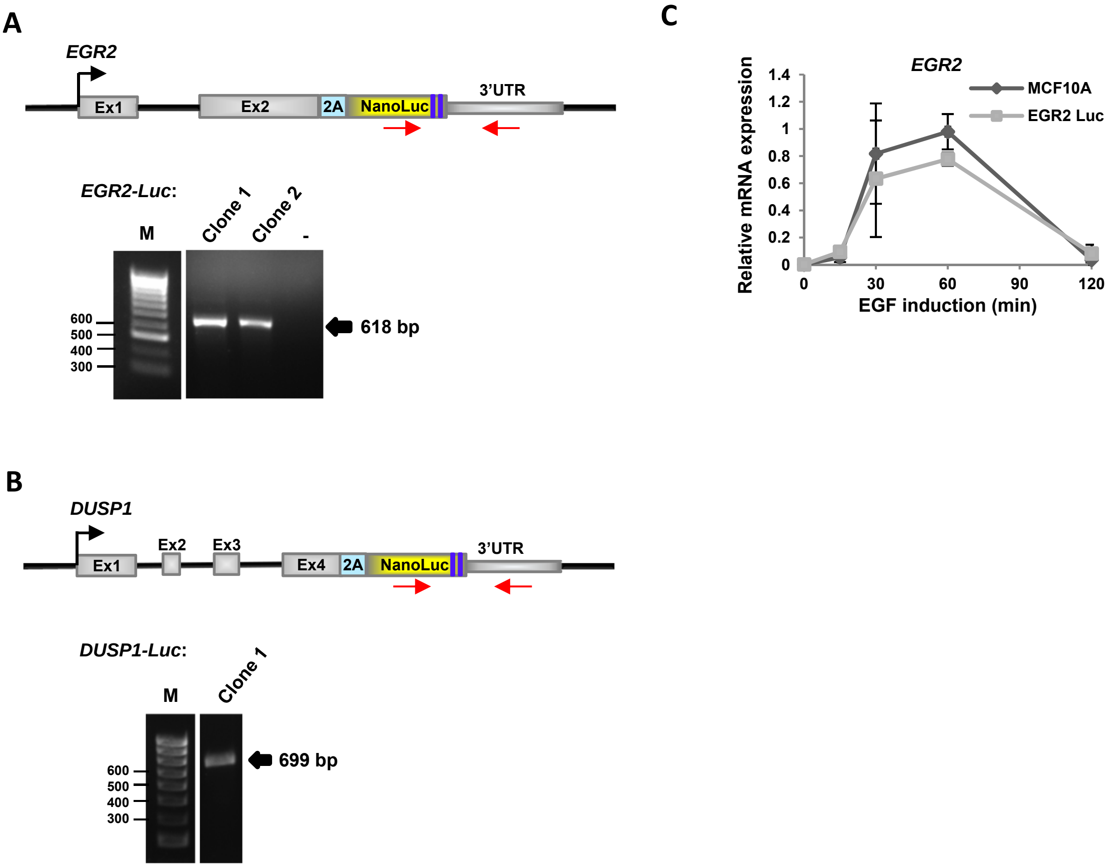


**Supplementary Fig. S1. Validation of the *EGR1*- and *DUSP1*-reporter systems.** (A and B) Diagrammatic representations of the newly created gene fusions (top) showing the locations of the PCR primers used for validation of the genomic insertion sites (one in the middle of nanoluciferase and one in the *EGR2* (A) or *DUSP1* (B) 3' UTR). PCR of genomic DNA (bottom) showing the expected sizes of product (618 bp and 699 bp) from positive clones. A control with no DNA added (-) is shown. (C) RT-qPCR analysis showing the relative mRNA expression of *EGR2* in Control (MCF10A) and MCF10A cells containing the *EGR2*-Luciferase reporter (EGR2-luc) following EGF addition. Similar induction profiles are seen in both cell lines.

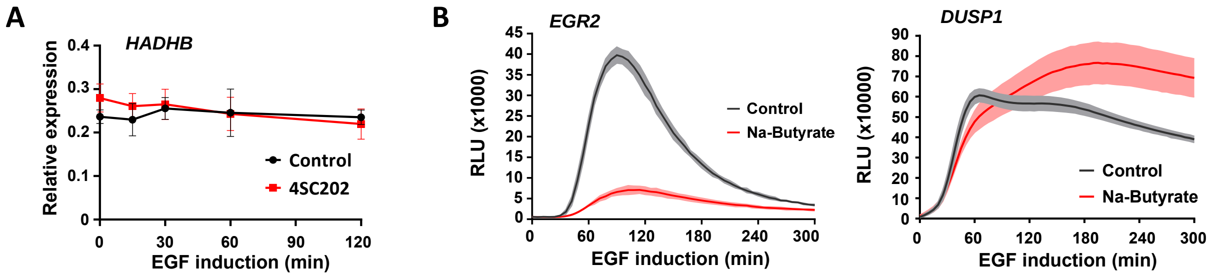


**Supplementary Fig. S2. Deacetylase inhibitors alter the *EGR2*- and *DUSP1*-reporter activation profiles.** (A) RT-qPCR analysis of control *HADHB* expression in MCF10A cells following EGF induction over 120 min in the presence and absence of the KDAC inhibitor 4SC202. (B) Activation kinetics of *EGR2*- and *DUSP1*-luciferase reporters after induction with EGF. Cells were treated with vehicle or Sodium (Na) butyrate 1 hr prior to EGF addition. Data are the average of three biological replicates (n=3), shaded area represents ±SEM.

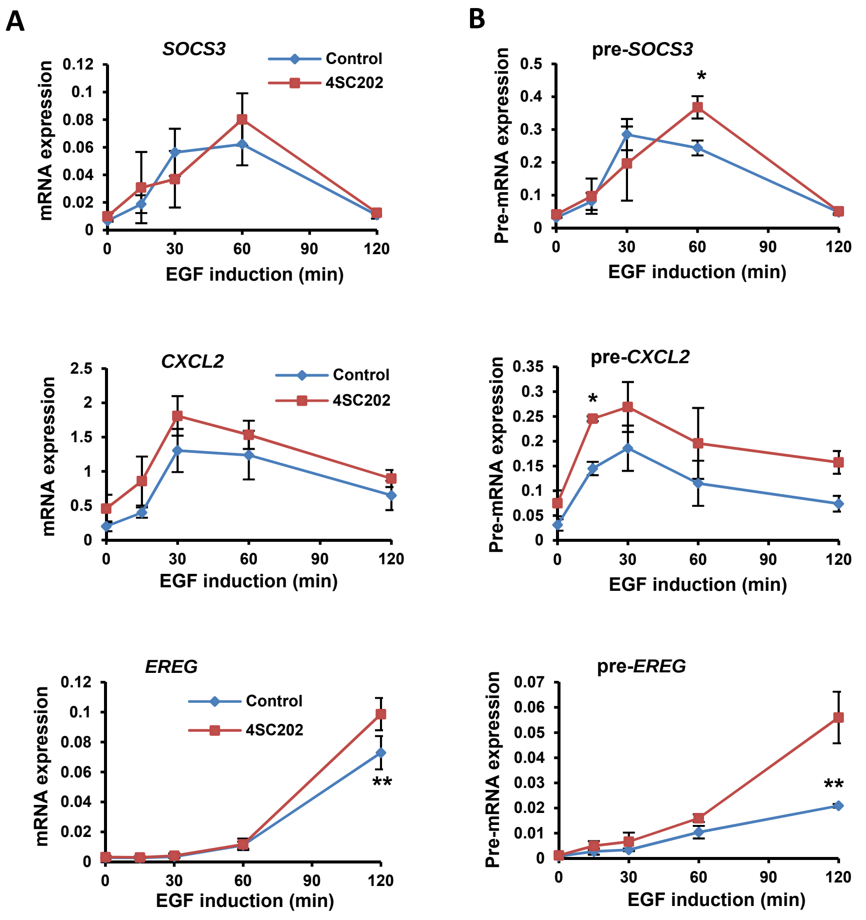


**Supplementary Fig. S3. Effect of deacetylase inhibition on EGF-mediated transcriptional activation**. Examples of relative mRNA (A) and pre-mRNA (B) expression profiles of genes from Figure 2C. One gene from group 1 (*SOCS3*) and two from group 2 (*CXCL2* and *EREG*) are shown after addition of EGF for the indicated times. Cells were treated with vehicle (Control) or 4SC202 for 1h before induction. Data represents mean of three biological replicates ± SD; *= P-value < 0.05, **= P-value <0.001.

A

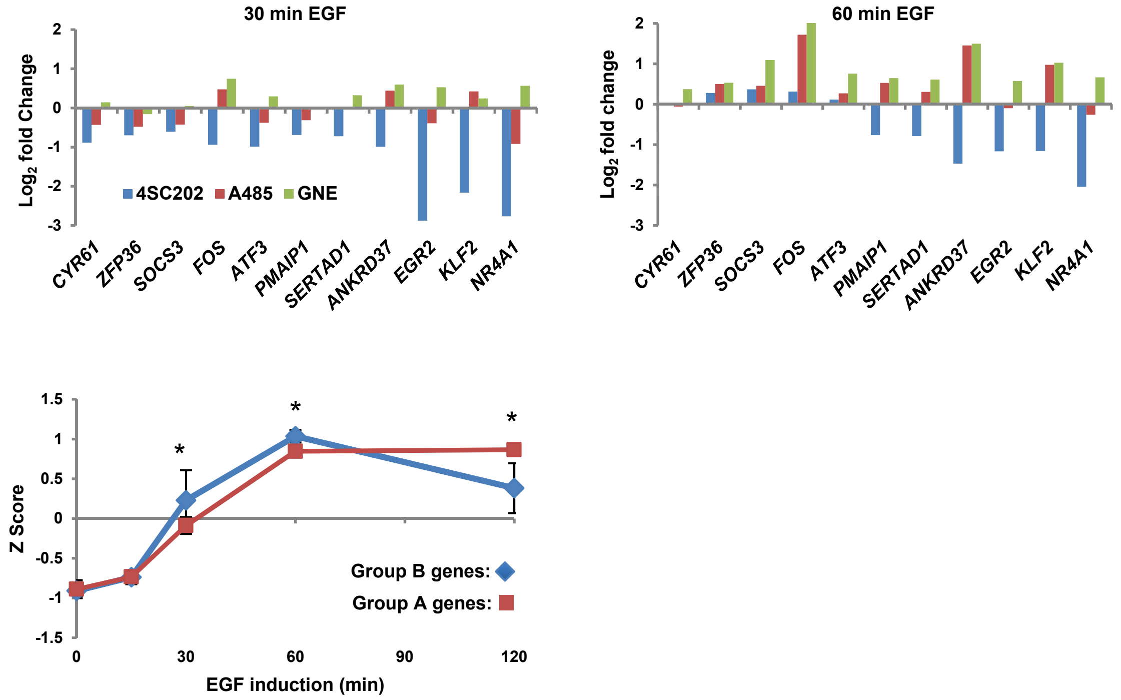


**Supplementary Fig. S4. Opposite gene expression responses to KDAC and KAT inhibition**. (A) Genes were selected from group B in figure 3B. Data are shown as log$^2$ fold change of expression after EGF addition for 30 and 60 min after pre-treatment with 4SC202, A485 and GNE-781. HAT inhibitors cause a general increase and HDAC inhibitors a general decrease in gene expression. (B) Z Score of mRNA expression of genes in group B (Fig. 3C and S4A above; blue line) and the genes in group A (Fig. 3C; red line) after induction with EGF at the indicated times. Lines represent means of Z scores ±SD; *= P-value <0.001.

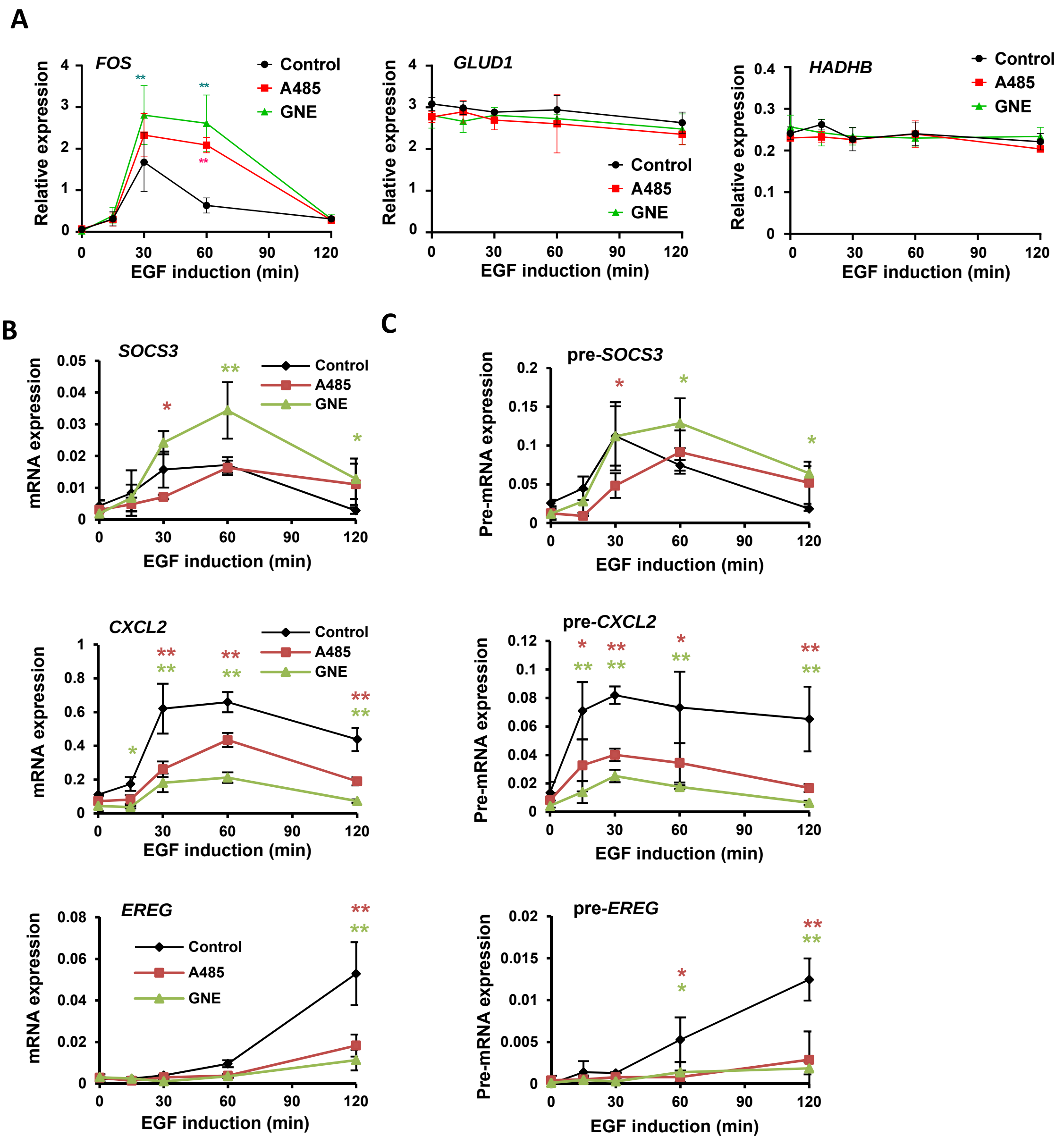


**Supplementary Fig. S5. Effects of KAT inhibitors on EGF-mediated gene activation profiles**. RT-qPCR analysis of gene expression in MCF10A cells following EGF induction over 120 min in the presence and absence of prior A485 or GNE-781 treatment. (A) Expression of *FOS* and the housekeeping genes *GLUD1* and *HADHB*. Relative mRNA (B) and pre-mRNA (C) expression profiles of the indicated genes after addition of EGF for the indicated times. One gene from group 1 (*SOCS3*) and two from group 2 (*CXCL2* and *EREG*) are shown. Data represents means of three biological replicates ± SD; *= P-value < 0.05, **= P-value <0.001. The effects on mRNA levels are mirrored by changes to pre-mRNA levels.

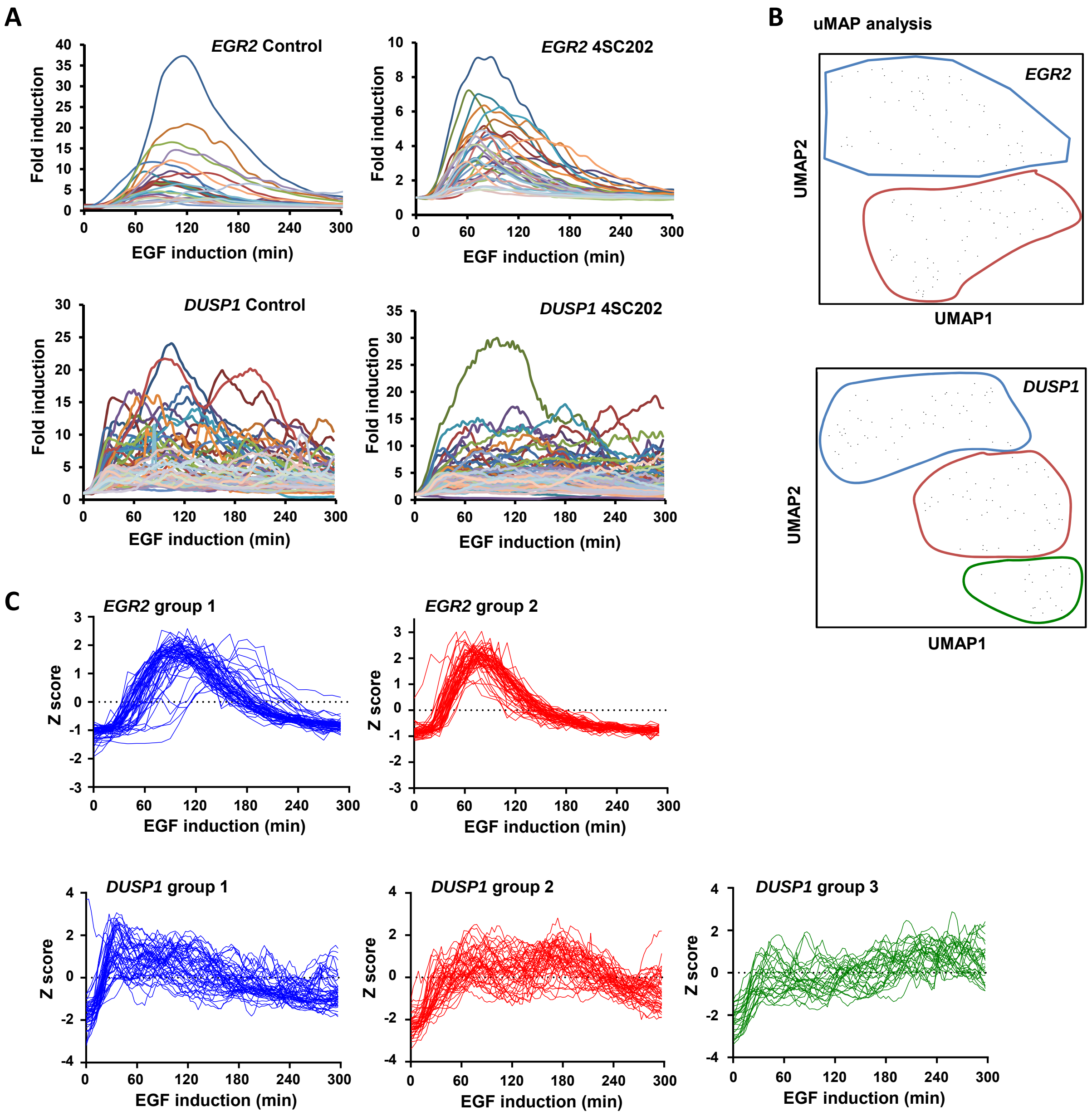


**Supplementary Fig S6. Analysis of single cell luciferase reporter profiles**. (A) Individual traces of *EGR2*- (top) and *DUSP1*- (bottom) reporter expression measured in single cells on the bioluminescence microscope. Cells were treated with vehicle or 4SC202 for 1h prior to EGF addition. Graph represents fold induction relative to zero time point (taken as 1). (B) Uniform Manifold Approximation and Projection (uMAP) analysis of 15 PCA groups for *EGR2*- and 10 PCA groups for *DUSP1*-reporter expression profiles. Enclosed are cells that cluster into different groups. (C) Kinetics of *EGR2*- and *DUSP1*-reporter expression in individual cells divided into different PCA defined groups (B). Each line the represents measured luciferase activity in one cell after EGF induction.

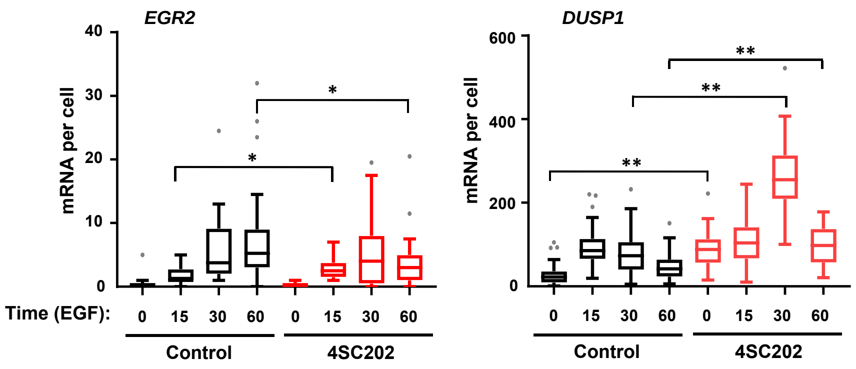


**Supplementary Fig S7. smFISH analysis pf *EGR2* and *DUSP1* expression profiles**. The numbers of mature mRNA transcripts of *EGR2* and *DUSP1* per cell measured by smFISH are shown as box plots with whiskers; horizontal lines represent median whiskers calculated by the Tukey method. Significance compared to control *= P-value < 0.05, **= P-value <0.001.
